# Adaptive stress response induced by toluene increases *Sporothrix schenckii* virulence and host immune response

**DOI:** 10.1101/539775

**Authors:** Damiana Téllez-Martínez, Alexander Batista-Duharte, Vinicius Paschoalini Silva, Deivys Portuondo Fuentes, Lucas Souza Ferreira, Marisa Campos Polesi, Caroline Barcelós Costa, Iracilda Zeppone Carlos

## Abstract

Environmental factors modify the physiology of microorganisms, allowing their survival in extreme conditions. However, the influence of chemical contaminants on fungal virulence has been little studied. Sporotrichosis is an emergent fungal disease caused by *Sporothrix schenckii,* a soil-inhabiting fungus that has been found in polluted environments. Here, we evaluated the adaptive stress response of *S. schenckii* induced by toluene, a key soil contaminant. The effect on fungal virulence and host immune response was also assessed. The fungus survived up to 0.10% toluene in liquid medium. Greater production of melanosomes and enhanced activity superoxide dismutase, associated to increased tolerance to H_2_O_2_ were observed in toluene-exposed fungi. Intraperitoneal infection of mice with *S. schenckii* treated with either 0, 0.01 or 0.1% of toluene, resulted in greater fungal burden at day 7 post-infection in spleen and liver in the groups infected with fungus treated with toluene 0.1%. A higher production of Il-1β, TNF-α, IL-10 and nitric oxyde by peritoneal macrophages and IFNγ, IL-4 and IL-17 by splenocytes was also observed in that group. Our findings showed that morphological and functional changes induced by toluene leads to increased *S. schenckii* virulence and antifungal host immune response in our model.

## Introduction

The impact of environmental factors on the emergence or outbreak of infectious diseases has become a major concern in public health (Wilson 1995; Giraud *et al*., 2010; Casadevall *et al*., 2003). Several lines of evidence show that acquired microbial resistance to extreme environmental factors can also enhance their resistance to host immune mechanisms, leading to increasing virulence (Casadevall *et al*., 2003; Casadevall and Pirofski 2007). Sporotrichosis is a worldwide subcutaneous mycosis caused by thermodimorphic fungi of the genus *Sporothrix* that contain several pathogenic species including *S. brasiliensis, S. schenckii sensu stricto, S. globosa, and S. luriei.* (Marimon *et al*., 2007; Rodriguez *et al*., 2018). These fungi are naturally found in soils plentiful in cellulose, with a pH range of 3.5 to 9.4, a mean temperature of 31 °C, and a relative humidity above 92% (Téllez *et al*., 2014). People acquire sporotrichosis by inoculation of fungal conidia in the environment after a traumatic lesion and more rarely by inhalation and mucosal contact (Barros *et al*., 2011). Currently, zoonotic transmission, mainly by cat scratch, is becoming a health problem in Brazil (Gremião *et al*., 2017). In recent decades, sporotrichosis has transformed from a rare disease to an emergent mycosis with an alarming growing incidence in several tropical and subtropical (Carlos and Batista-Duharte 2015; Chakrabarti *et al*., 2014).

Sporotrichosis outbreaks have been described in large areas of poverty with a high demographic density and precarious sanitary and health system (Barros *et al*., 2008; Silva *et al*., 2012). *S. schenckii* species has been isolated in highly polluted environments with chemical contaminants. (Dixon *et al*., 1991; Ulfig 1994; Ulfig and Lukasik 1996; Kusuhara 2009; Pečiulytė 2010). Several studies have also shown that *S. schenckii* is able to resist the effect of radiation (Torres-Guerrero and Arenas-López 1998; de Souza Lacerda *et al*., 2011), very low temperatures (Pasarell and McGinnis 1992; Mendoza *et al*., 2005; Ramírez-Soto *et al*., 2018.), high osmotic pressure for several years (Borba and Oliveira 1992; de Capriles *et al*., 1993; Mendoza *et al*., 2005) and high relative humidity (Ramírez-Soto *et al*., 2018.). These findings suggest a robust ability of *S. schenckii* to resist extreme environmental conditions (Téllez *et al*., 2014; Batista-Duharte et al. 2015).

A recent study of our group revealed that mice exposed to mercury (II) chloride enhances susceptibility to *S. schenckii sensu stricto* infection (Batista-Duharte *et al*., 2018). However, there are not studies on the effects of chemicals pollutants on *Sporothrix spp* biology (Téllez *et al*., 2014; Ramírez-Soto *et al*., 2018).

Aromatic compounds such as benzene, toluene, ethylbenzene and xylene isomers, collectively known as BTEX, are one of the major contributors to environmental pollution. BTEX are extensively used as solvents in many industrial processes and as base reagents for the production of chemicals products. In addition, BTEX are components of gasoline and aviation fuels. They are often released in the environment during production, transport, use and disposal, causing groundwater, surface water, and soil contamination. Toluene has been found in at least 1,012 of the 1,699 National Priorities List sites identified by the Environmental Protection Agency (EPA) and it is included in the Agency for Toxic Substances and Disease Registry (ATSDR) 2017 substance priority list, which is based on a combination of their frequency, toxicity, and potential for human exposure (*Agency for Toxic Substances and Disease Registry. ATSDR’s Substance Priority List.* 2017).

Prenafreta-Boldu isolated several fungi, including *S. schenckii*, in biofilters exposed to polluted air or hydrocarbon gas streams. The data from this research showed that volatile hydrocarbon-degrading strains are closely related to a very restricted number of pathogenic fungal species that cause severe mycoses, especially neurological infections, in immunocompetent individuals (Prenafeta-Boldú *et al*., 2006).

To date, there have been no studies examining the influence of chemical contaminants on the biology of *S. schenckii sensu latu.* Given that this fungus inhabits soils and this niche is frequently contaminated with oil and their products, the objective of this work was to evaluate the resistance of this fungus to toluene and it influence on fungal virulence. The results of this work can help to understanding how chemical contamination can influence fungal virulence and possibly certain sporotrichosis outbreaks in contaminated regions.

## Materials and Methods

### Microorganisms and preparations

The *S. schenckii sensu stricto* strain ATCC 16345 (here named as *S. schenckii*) used in this experiment was originally obtained from a human case of diffuse lung infection (Baltimore, MD) and kindly provided by the Oswaldo Cruz Foundation (Rio de Janeiro, Brazil). The mycelial phase was maintained at room temperature in Mycosel (BD Biosciences) agar slants. A piece of the fungal mycelium (approximately 1 cm^2^) was grown in 100 mL of Sabouraud Dextrose Broth (SDB) (Difco, Detroit, USA) for 4 days on a rotatory shaker at 130 rpm and 30°C. The conidia were then isolated from the hyphae by filtration through several layers of sterile gauze using a Buchner funnel and suspended in 1 mL of phosphate-buffered saline (PBS). After incubation, conidia aliquots were <u>seeded</u> for the different experiments in SDB medium and then incubated for 5 days under the same conditions described above.

### Effect of toluene concentration on S. schenckii growth

Tests were performed in 125-mL Erlenmeyer flasks containing 50 mL of SDB and sealed with Teflon Mininert valves (SUPELCO, 24 mm) to prevent evaporation of the solvent. SDB was supplemented with toluene at different concentrations (0.01, 0.1, 1.0 e 10.0% (vol/vol)). An aliquot of 1×10^7^ conidia was inoculated and incubated at 30°C and 130 rpm on a rotatory shaker. Control cultures without toluene were included. Fungal survival was determined after 24 hours of incubation by counting Colony Forming Units (CFU) on Petri dishes containing Sabouraud Dextrose Agar (SDA). Percent survival was determined by comparing the number of colonies of fungal cell grown with toluene to those grown non-toluene.

### Fungal growth in toluene

The growth of *S. schenckii* in two concentrations of toluene was evaluated according to the previous results and under the same conditions. The fungal growth was monitored during five days by CFU count. The LIVE/DEAD® Yeast Viability Kit (Molecular Probes) dye test was performed in accordance with the manufacturer’s instructions, to evaluate the cell viability during the exponential phase of the fungal cultures. Samples were visualized on the BH50 fluorescence microscope (Olympus) using the fluorescein and DAPI filters for the FUN1 and Calcofluor White M2R stains, respectively.

### Toluene consumption by fungal cells

Cultures supplemented with 0.01% (v/v) toluene were used to evaluate the growth and consumption of solvent. Tests were performed as previously described. The concentration of toluene in the cultures was quantified by autosampler injection in headspace on a gas chromatograph (GC) (Perkin Elmer-Clarus 680) equipped with a flame ionization detector (FID). A sample was injected into an Elite-624 column (30 m x 0.25 mm x 1.4 µm; 6%-cyanopropylphenyl-94% dimethylpolysiloxane). Helium served as the carrier gas (1 mL/min) and hydrogen as the fuel gas for FID. Residual toluene was quantified by comparison of the area under of the curve determined by chromatography with those obtained for the curves standards. Samples were collected every 24 hours and analyzed under the same conditions as the standards.

### Transmission Electron Microscopy (TEM)

Conidia grown for 5 days in 0; 0.01 and 0.10% toluene were studied by TEM to evaluate morphological modifications. Conidia were incubated overnight at 4°C in fixation solution containing 2.5% glutaraldehyde and 2% paraformaldehyde in 0.1 M cacodylate buffer (pH 7.2) (Sigma-Aldrich). The cells were then washed three times, exposed to potassium dichromate and stained with 0.5% uranyl acetate overnight at 4 °C. The samples were washed, soaked in agar and cut into small cubes of approximately 0.5 to 1 mm^3^, and they were dehydrated in increasing concentrations of ethanol, placed in Spurr resin, sectioned in ultrafine blocks with an ultramicrotome and placed in copper grids. The observations were performed using a Jeol JEM-100 CXII transmission electron microscope equipped with a Hamamatsu ORCA-HR digital camera. Digital images were captured at 5,000X and 40,000X. The thickness of the cell wall was determined using four independent measures for each image, from the random placement of a grid of cardinal points (Renzoni *et al*., 2011). The reported values represent the mean of 42 cells for each group. Measurements were determined using ImageJ software. The conidial area was measured using ImageJ software (National Institutes of Health, Bethesda, MD USA).

### SOD activity assay

**S**OD activity was photochemically assayed on 10% polyacrylamide gels (NATIVA-PAGE) based on the nitro blue tetrazolium (NBT) photoreaction and in Tris-glycine buffer under non-denaturing conditions according (Kuo *et al*.,. 2013; Weydert and Cullen 2010).

The SOD enzyme extract was prepared according to Niyomploy *et al*., (2014) with some modifications. Fungal cells were cultivated with 0.01 and 0.10% (v/v) of toluene as described above. Cell cultures were centrifuged at 500xg for 10 minutes and washed three times with ice cold 25 mM Tris-HCl buffer (pH 8.5). Five grams of fungal mass from each treatment were sonicated with 25 mM Tris/HCl extraction buffer (pH 8.5), containing 2 mM DTT (Sigma-Aldrich, St. Louis, USA), and 5 mM ethylenediamine tetraacetic acid (EDTA) (Sigma-Aldrich, St. Louis, USA), and 1 mM PMSF to prevent proteolytic activity. The supernatants were collected after centrifugation and then subjected to 80% ammonium sulfate (NH4)_2_SO_4_. The mixtures were incubated overnight, and then centrifuged at 17880xg for 30 minutes. The precipitates were resuspended, dialysed with deionized water and concentrated in tubes AMICON Ultra-3k (Merck-Millipore) according to the manufacturer’s instructions. Concentrated volumes were collected and quantified. The samples were stored at −80 °C until separation in electrophoresis to determine the enzymatic activity. The entire extraction and purification process was carried out at 4 ° C.

### Sensitivity to oxidative stress

Aliquots of 2×10^7^ conidia/mL grown for 5 days in 0; 0.01 and 0.10% toluene were suspended in sterile deionized water. Exponential dilutions of this aliquot were performed in 96-well plates containing the different concentrations of hydrogen peroxide (H_2_O_2_) (between 0 and 125 mM) and incubated at 28 °C for 90 minutes under mild agitation. Then, each dilution was seeded in Petri dish containing SDA and incubated at room temperature during 6 days. The addition of fungus without toluene served as a control (Ramírez-Quijas *et al*., 2015).

### Measurement of Intracellular Reactive Oxygen Species (ROS)

The ROS-specific dye DHR-123 is oxidized to the fluorescent rhodamine 123 by the intracellular ROS that could be detected by flow cytometry. Endogenous ROS was measured as previously described (Shao *et al.* 2016) with modifications using the fluorescent dye dihydrorhodamine-123 (DHR-123, Sigma–Aldrich, USA). The results were recorded and analyzed using a FACS Accuri C6 flow cytometer (Becton-Dickinson, USA).

### Animals and experimental design

To determine whether changes induced by toluene in the in fungal cells caused modifications in their virulence, we used an experimental model of systemic *S. schenckii* infection described by Ferreira *et al*., 2015. Male BALB/c mice, 5–7 weeks old at the time of inoculation, were purchased from “Centro Multidisciplinar para Investigacão Biológica na Área da Ciência de Animais de Laboratório” (CEMIB), UNICAMP University (Brazil). Animals were housed in individually ventilated cages in an ambient with controlled temperature and 12-h light/dark cycles. Water and food were offered ad libitum. Animals were intraperitoneally inoculated with 10^6^ *S. schenckii* conidia cells grown with toluene or at either 0.01 or 0.1% toluene concentration. For inoculation conidia were suspended in PBS (10^6^ conidia/mL). A control non-infected group was inoculated with an equal volume of PBS alone. Assessment of the systemic fungal load was performed at day 7th post-infection by counting the CFU grown on Mycosel agar plates of liver and spleen macerates. All animal procedures were approved by the Research Ethics Committee of the Faculty of Pharmaceutical Sciences of Araraquara – Estadual Paulista University (UNESP) (Protocol nº 48/2015) and were in accordance with the National Institutes of Health Animal Care guidelines and the guidelines of the Brazilian College of Animal Experimentation (COBEA).

### Measurement of the ex vivo release of IL-1β, TNFα, IL-10 and nitric oxide (NO) by peritoneal macrophages

Thioglycollate-elicited peritoneal exudate cells (PECs) were harvested from male Balb/c mice 3 days after i.p. inoculation with 3% sodium-thioglycollate, which was performed by washing the peritoneal cavity with 5.0 mL of sterile PBS, pH 7.4. The cells were cleaned twice by centrifugation at 200g for 5 min at 4 °C and washing with sterile PBS, pH 7.4. The cells were then resuspended in RPMI-1640 medium (Sigma) supplemented with 100 U/mL penicillin, 100 μg/mL streptomycin, 5 × 10-2 M mercaptoethanol, and 5% inactivated fetal calf serum (Sigma), referred to as RPMI-1640 complete (RPMI-1640C)medium. PECs were manually counted in a Neubauer chamber (Boeco, Germany), and the concentration was adjusted to 5 × 106 PEC/mL in RPMI-1640C for use in the following experiments. 0.1 mL PEC suspensions containing 5 × 106 cells/mL in RPMI-1640C were added to each well of a 96-well tissue culture plate and incubated for 1 h at 37 °C with a supply of 5% CO2. Next, non-adherent cells were removed by discarding the supernatants and refilling each well with 0.1 mL of RPMI-1640C medium (Batista-Duharte *et al*., 2016).

Peritoneal macrophages were treated with HK conidia (HKC) from each group or lipopolysaccharide (LPS) as positive control `for` 24 h in 5% CO2 at 37°C in corning costar flat bottom cell culture plates 24-well. The cell-free supernatants were assayed for IL-1β and TNFα using a commercial Enzyme-Linked Immunosorbent Assay (ELISA) Kit (BD Bioscience-Pharmingen) according to the manufacturer’s instructions. Colorimetric reactions were measured at 450 nm with wavelength correction at 570 nm (Microplate Reader, Multiskan Ascent, Labsystems).

For the evaluation of NO content, the Griess reaction was used. Briefly, 100 μL of supernatant was incubated with 100 μL of Griess reagent (1% sulfanilamide (Sigma Chemicals, St. Louis, MO, USA), 0.1% N-(1-naphthyl)-ethylenediaminedihydrochloride (NED, Sigma Chemicals, St. Louis, MO, USA) and 2.5% phosphoric acid in water). The NO concentration was determined using the same microplate reader at 540 nm, with reference to a standard curve (NaNO_2_)

### Measurement of the ex vivo release of IFN-γ, IL-17 and IL-4 in splenocytes

Spleens were aseptically removed and passed through a 100_m cell strainer into a Petri dish containing 2mL of PBS with the aid of a syringe plunger. For red cell lysis, the resulting suspension was added with 6mL of a 0.17Mammonium chloride solution and then incubated on ice for 5 min. The splenocytes were then separated from the supernatant by centrifugation at 300×g for 5 min at 4 ?C, washed once with 3mL of RPMI complete medium and then resuspended in 1mL of the same medium. Cell concentration was determined by microscopy using the Trypan blue exclusion test and then the splenocytes were adjusted to 5×10^6^ cells/mL in RPMI complete medium. Adjusted splenocytes were treated with HK conidia (HKC) from each group or Concanavalin A (ConA) as positive control `for` 24 h in 5% CO2 at 37°C in corning costar flat bottom cell culture plates 24-well. The cell-free supernatants were assayed for IFN-γ, IL-17 and IL-4 using a commercial ELISA Kit (BD Bioscience-Pharmingen) according to the manufacturer’s instructions

### Statistical analysis

All data was performed using GraphPad Prism version 6.01. The quantitative data in the figures are presented as the mean +/-standard error of the mean. A one-way or two-way analysis of variance (ANOVA) followed by Tukey’s method for comparison of multiple groups was used. All the experiment were performed in triplicate in independent assays. For all experiments, a statistical comparison was considered significant when p < 0.05 as indicated on the figures with asterisks.

## Results and discussion

### Effect of toluene concentration on S. schenckii

The fungal cell concentration (CFU) drastically decreased when grown to 0.01% toluene by approximately 80% in 24 hours, while 0.10 caused more than 90% mortality. Higher concentrations were lethal for 100% of the seeded conidia (Figure 1 A). The (CFU) decreased 0.38 log units, while at 0.10% it was reduced 1,394 units when compared to the fungus growing without toluene.

**Figure 1.**
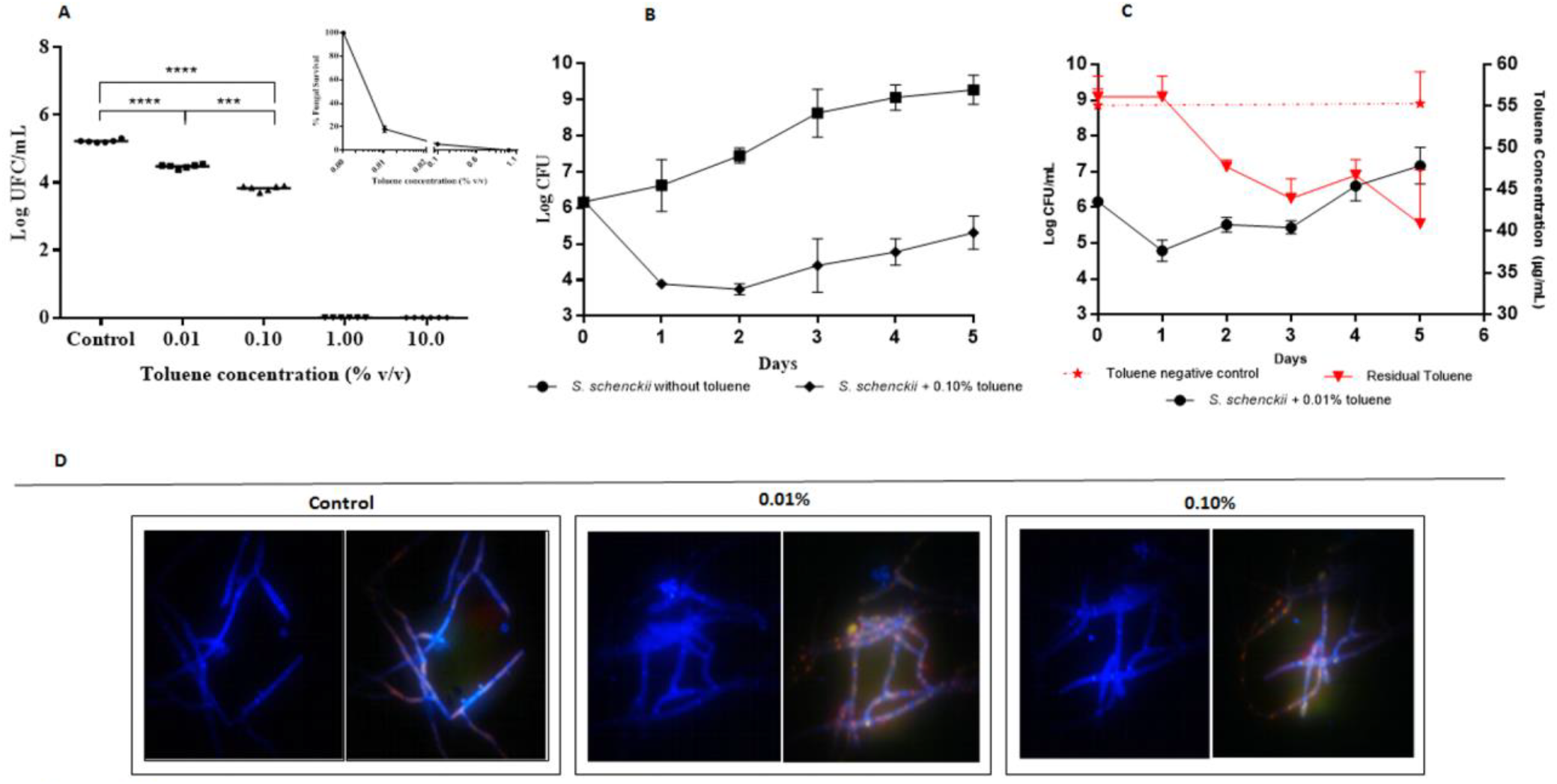
A) Survival of *S. schenckii* cultured in Sabouraud dextrose broth in the absence or presence of toluene at the indicated concentration. Colony forming units (CFU)/mL expressed in logarithmic notation. Percent survival compared with non-treated controls. ***(p <0.001): ****(p <0.0001). B) Growth kinetics of *S. schenckii* without toluene (black line) or in the presence of 0.01% toluene (v/v) (continuous red line) under aerobic conditions and a constant temperature of 30°C. Discontinuous red line: concentration of toluene in sterile flasks. Blue line: residual toluene in culture with growing fungus. The toluene concentrations were plotted as the mean ± standard deviation of two independent experiments. C) Calcofluor white-stained *S. schenckii* showing cell wall integrity (left pane) and FUN1®-stained cells showing orange intravacuolar fluorescence cell viability (right) of live fungi in the mycelial phase.

### Growth in toluene. Consumption by fungal cells

All cultures started from a viable cell concentration of 1,5×10^6^ CFU/mL. Both concentrations of toluene produced a reduction of the population at the first 24 hours. At concentration of 0.01% toluene the fungal population was reduced between 1.5 and 2 log units, to about 6.13×10^4^ CFU/mL. After the latency phase the fungus increased the CFU and at 5 days of incubation reached a concentration of 1.47×10^7^ CFU/mL on average. At concentration of 0.01% toluene the fungal population was reduced from 6.21 to 3.89 log units (7.76×10^3^ CFU/mL), completing the latency phase of 48 hours followed by a gradual increase in cell concentration, which reached 2.06×10^5^ CFU/mL. (Figure 1 B, C). The viability and cell wall integrity of the fungi were checked on the fifth day by Calcofluor and Fungal V staining, respectively. The tests confirmed the presence of structurally integrated and metabolically active mycelia in control and treated fungi (Figure 1 D).

Moreover, the consumption of toluene by growing *S. schenckii* was evaluated at 0.01% for 5 days and a culture free of toluene was used as control. As shown in Figure 1C, during the first 24 hours toluene concentration in the culture remained unchanged. After this interval, the solvent decreases in correspondence with the beginning of the exponential growth phase of the fungus. Approximately 26% of the toluene supplied to the study system was consumed at 72 hours after initiation of incubation. A continuous reduction in toluene concentration was observed as cell concentration increased. This result suggest that *S. schenckii* was able to metabolize toluene, as previously described (Prenafeta-Boldú *et al*., 2006).

### Conidial morphological changes

Figure 2 shows the ultrastructural changes in non-treated and toluene-treated fungal conidia under transmission microscopy. Conidia from fungi exposed to toluene showed a higher number of black-colored electrodense bodies (melanosomes) in the cytoplasm and a reduction in the thickness of the cell wall compared to non-exposed conidia (Figure 2A,B,C). The number of conidia with melanosomes was also higher in conidia grown in toluene compared with non-exposed conidia Figure 2D). Conidia with low-density electron structures in the periphery and high-density structures in the center were also observed (Figure 2A), which appeared to be melanosomes in the early stages of formation (Upadhyay *et al*., 2016).

**Figure 2.**
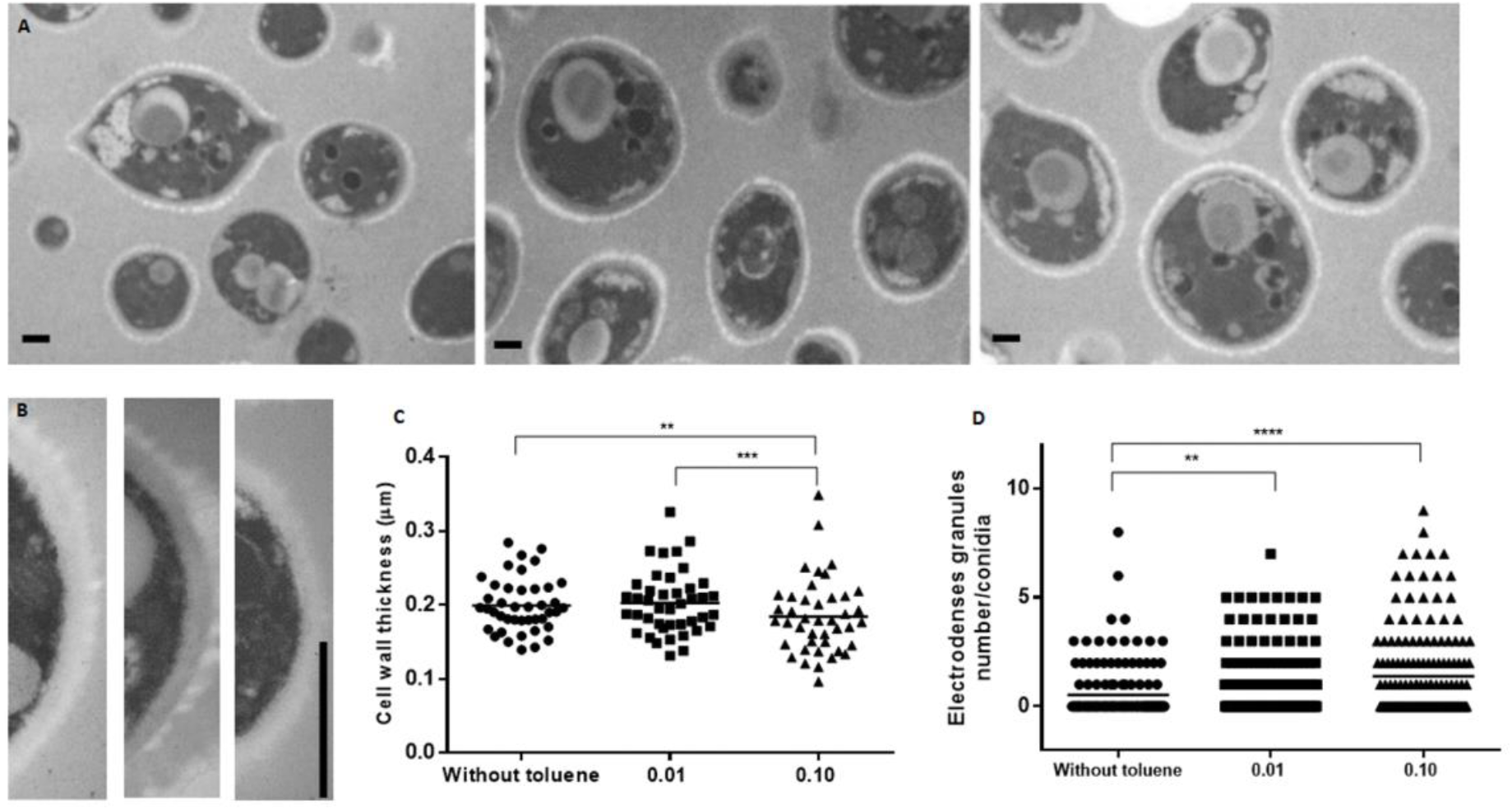
Representative transmission electron microscopy (TEM) images of conidial cells from non-treated or 0.01 or 0.1% toluene-treated *S. schenckii*. A) Accumulation of electrodense material in cytoplasmic vesicles (melanosomes) of *S. schenckii* conidia exposed to toluene. B) Fungal cell wall thickness. C) Scattergram of cytoplasmic melanosome count and D) fungal cell wall thickness. Ultrafine sections of fungal cells were prepared for TEM as previously described. The data represent the mean ± standard deviation of 161 conidia in each group (magnification 5000x, 40.000X, bar = 1 μm) ****(p<0.0001).

Melanin is a virulence factor that enhances fungal resistance to environmental stressors and immune responses in the host (Eisenman and Casadevall 2012). The natural presence of melanin has been demonstrated in *S. schenckii* species and it favors the fungal virulence and resistance to antifungal drugs (Morris-Jones *et al*., 2003; Madrid *et al*., 2011; Almeida-Paes *et al*., 2016). Here, increased production of cytoplasmic melanosomes in different phases of formation was observed in conidia of fungi exposed to toluene, especially at the highest concentration. Different reports have revealed that fungal melanization occurs in specialized vesicles that are analogous to mammalian melanosomes. Melanization in vesicles in form of melanosomes is necessary because the reaction generates different highly reactive intermediates that self-react to create melanin. Thus, melanosomes can protect the cells from the toxic effects of these metabolites (Nosanchuk *et al*., 2015). Internal melanosomes or melanosome-like structures has been observed in *Fonsecaea pedrosoi* (Alviano *et al*., 1991; Franzen *et al*., 1999; 2008). *C. albicans* (Walker *et al*., 2010), Cladosporium carrionii, Hormoconis resinae (San-Blas *et al*., 1996).

Garrison *et al*., (1979) reported for the first time the presence of electro opaque bodies (EOB) in the cytoplasm of *S. schenckii* yeasts contained in biopsy material obtained from naturally-occurring disseminated feline sporotrichosis. The inclusions described were analogous to EOB previously reported in *S. schenckii* hyphae `by the same group` (Garrison et al., 1977). Although the EOBs described by Garrison and coworkers were structurally similar to the later described melanosomes-like structures, these authors did not relate these structures to melanin formation. This function was proposal was recently proposal by Almeida-Paes *et al*., (2017) in *S. schenckii* yeast. In our study, the stimulation of melanosomes production under fungal stress in *S. schenckii* conidia is reported for the first time.

The expression of SOD was evaluated by non-denaturing electrophoresis. In the gel, SOD was clearly identified as achromatic bands owing to the inhibition of NBT. Inhibition of NBT is a measure of the amount of SOD present in the sample and shows the detoxification capacity of the fungus in the presence of toluene. In the control sample (without toluene), there was low expression of SOD, at least at the tested levels (Figure 3C). Qualitatively, the production of catalase was observed by the immediate effect of H_2_O_2_ on colonies of *S. schenckii* that grew in the presence of toluene and non-treated fungi. In the first 15 seconds, fungal colonies that grew in the presence of 0.1% toluene exhibited more profuse and prolonged bubble formation than both the control group and those grown in the presence of 0.01% toluene (Figure 3D).

**Figure 3.**
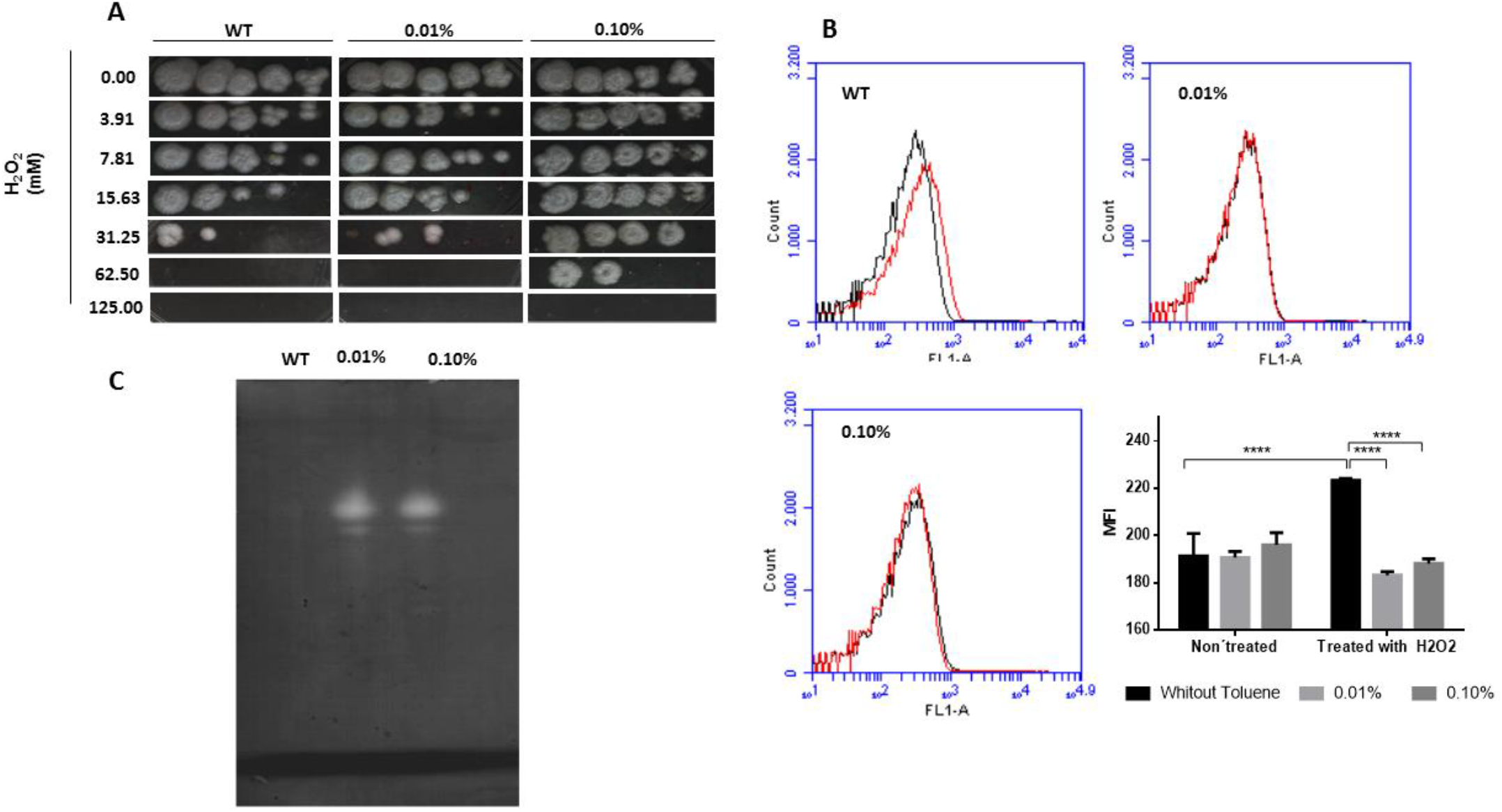
Antioxidant mechanisms. A) *S. schenckii* grown in toluene become more resistant to hydrogen peroxide stress. Enzymatic mechanisms for the detoxification of reactive oxygen species developed by fungus grown in toluene allowed it to survive up to oxidant concentrations of 62.50 mM. B) Changes in superoxide dismutase activity. (1): Without toluene; (2): 0.01% toluene; (3): 0.10% toluene. To each well was added 23 μg of protein. C) Catalase activity observed by the reaction force and evolution of bubble formation (oxygen). D) Accumulation of ROS in *S. schenckii* cells stained with DHR-123 and detected by flow cytometry. Conidia grown without and with toluene were treated with H2O2 (red line), and untreated cultures with H2O2 (black line) were used as controls. E) Graphic representation of the mean fluorescence intensity. The tests were performed in triplicate.

### S. schenckii grown in toluene becomes more resistant to stress by hydrogen peroxide

*R*eactive oxygen species (ROS) production increases in fungi due to various stress agents such as starvation, radiation, mechanical damage, and others deleterious effects (Gessler *et al.* 2007). The main mechanism responsible for the detoxification of ROS depends on enzymatic systems formed mainly by antioxidant enzymes such as catalase, superoxide dismutase, and glutathione peroxidase, among others (Angelova et al 2005). Fungal resistance to hydrogen peroxide (H_2_O_2_) depends on the effectiveness of their antioxidant system and it is associated to fungal virulence. Compared with *Saccharomyces cerevisiae*, which exhibits great sensitivity to hydrogen peroxide challenges, pathogenic *Candida albicans* has a high natural resistance to oxidant agents (Pedreño *et al*., 2006).

Studies have demonstrated that toluene triggers the expression of fungal antioxidants to induce cell detoxification and to reduce the impact of the damage caused by ROS (Blasi *et al*., 2017). In this study, fungi growing at 0.01 and 0.10% toluene showed marked resistance to H_2_O_2_. Those fungi that grew in the presence of 0.10% toluene showed more robust growth when compared to the fungi that were not grown to toluene (control). They were able to grow at double the concentration of H_2_O_2_; the non-treated fungi with toluene were resistant up to 31.25 mM of H_2_O_2_, while the fungus treated with 0.01% and 0.10% toluene were resistant up to 31.25 and 62.5 mM of H_2_O_2_, respectively (Figure 3A).

The intracellular ROS was measured with DHR-123 dye, which is oxidized to Rhodamine 123 by ROS, and then detected using a flow cytometer. Non-treated conidia with toluene produced more ROS that those treated with toluene after exposure to H_2_O_2_ (Figure 3B). As show in Figure 3A, D, the median fluorescence intensity (MFI) conidia non-grown in toluene and exposed to H_2_O_2_ presented high levels of ROS. Levels of intracellular ROS of conidia grown in toluene and exposed to H_2_O_2_ were lower when compared with those not treated with toluene but exposed to the oxidant (Figure 3B,C,D).

*Cladophialophora immunda* strains are environmental fungi that cause opportunistic subcutaneous phaeohyphomycosis in immunosuppressed patients (Badali *et al*., 2008). Members of this group show a special association with hydrocarbon-polluted environments and can grow in the presence of toluene as the sole carbon and energy source. Melanization and the presence of a powerful antioxidant system are two characteristics of this species that allow *C. immunda* to survive in contaminated environments and simultaneously participate in its pathogenicity (Blasi *et al*., 2016). Experimental exposure to toluene, similar to the present study, triggers *C. immunda* expression of antioxidants as well as genes involved in cell detoxification (Blasi *et al*., 2016).

### Adaptive stress response increases S. schenckii virulence

Our next step was to assess the fungal virulence and the immune host response against *S. schenckii* after exposed our non-exposed to toluene. As seen in figure 4, fungi exposed to toluene achieved greater colonization in liver and spleen manifested by a higher count of colony - forming units in both organs, mainly in animals infected with conidia treated at the highest concentration of toluene.

**Figure 4.**
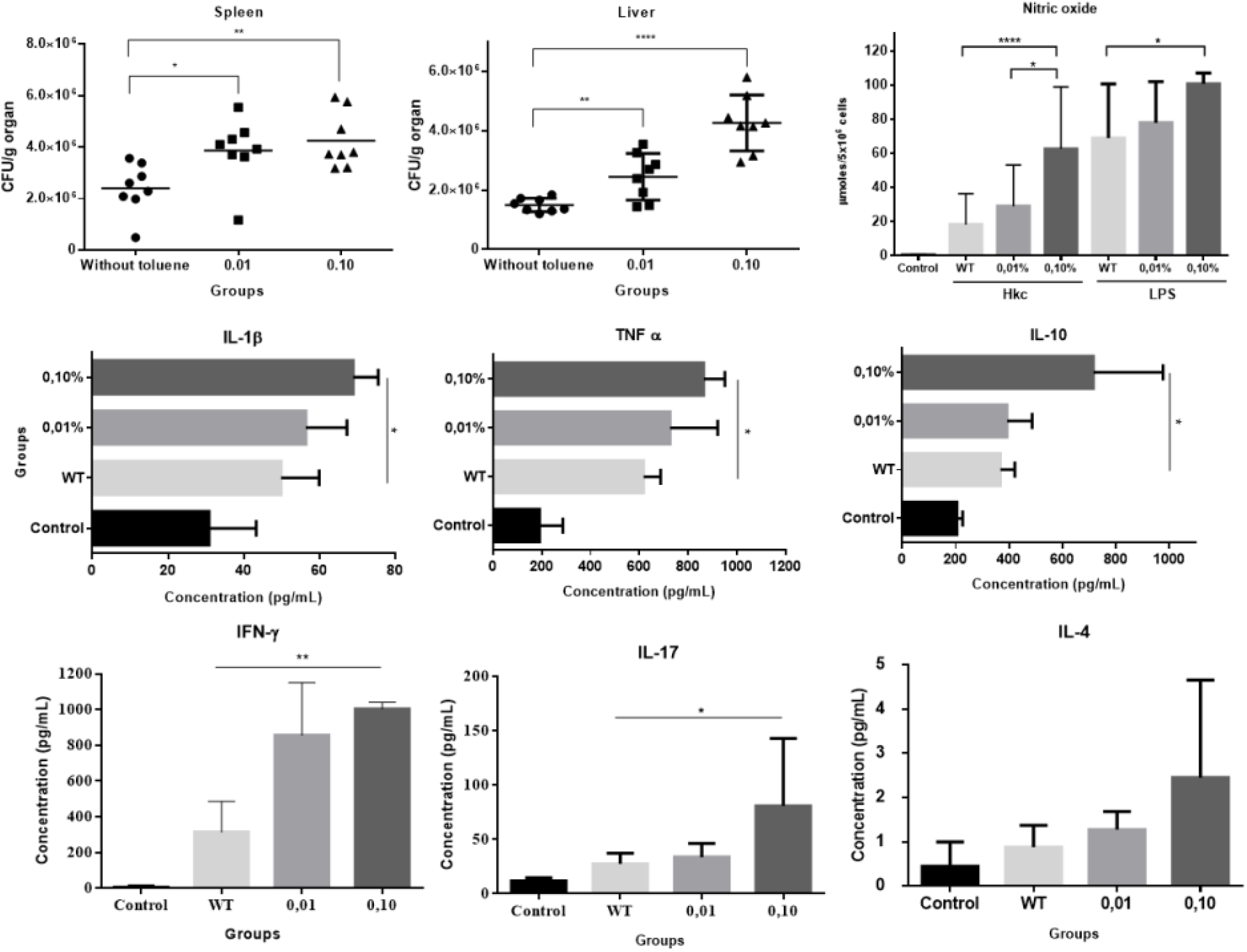
Scattergram of colony forming units (CFU)/g of spleen and liver from mice infected with either non-treated or 0.01 or 0.1% toluene-treated *S. schenckii.* Production of nitric oxide (NO), IL-1β, TNF-α and IL-10 by peritoneal macrophages, and IFN-γ, IL-17 and IL-4 by splenocytes. Cytokines and NO were evaluated in supernatant culture after overnight *in vitro* stimulation with heat killed conidia. Each animal was inoculated intraperitoneally with 1 x 10^6^ conidia, and infection was evaluated at day 7 post-infection. Data are shown as the mean ± SD of three independent experiments *(p <0.05); ** (p <0.01); ***(p <0.001); ****(p <0.0001).

In parallel, proinflammatory cytokines and nitric oxide produced by peritoneal macrophages in infected mice were quantified after *in vitro* stimulation with HKC conidia. The highest production of IL-1β, TNF-α, IL-10 and NO was observed in the group infected with 0.10%. A balanced production of the proinflammatory IL-1β, TNF-α and the antinflammatory cytokine IL-10 is estimulated during *S. schenckii* infection (Alegranci *et al*., 2013; Maia *et al*., 2016). However, although NO is also stimulated and it has an antifungal effect in vitro against this fungus, it also seems to have an immunosuppressive effect that favors fungal infection (Fernandes *et al*., 2008; Gonzalves *et al*., 2015).

The response of IFN-γ, IL-4 and IL-17 triggered by splenocytes from infected mice were also assessed after *in vitro* stimulation with HKC. These cytokines were quantified in the culture supernatants to evaluate the T helper cell type 1 (Th1), Th2 and Th17 responses respectively. A greater response of all these mediators was observed in the animals infected with the treated fungi, principally those that grew at the highest concentration of toluene (Fig 4). Several studies reveals that a Th1 and Th17 are stimulated during *S. schenckii* infection and they are important for the fungal clearance (Ferreira *et al*., 2015; Gonçalves *et al*., 2017; Batista-Duharte *et al*., 2018). Interestingly, a recent report of our group reveled that both *S. schenckii* and the more virulent *S. brasiliensis* stimulate similar Th1 response, but *S. brasiliensis* induced a more severe disease associated with sustained Th17 and a regulatory T cells (Tregs) responses than *S.* schenckii (Batista-Duharte *et al*., 2018). We are developing new analyzes to deepen the immunological mechanisms associated with the greater virulence of the group infected with the fungus exposed to 0.1% of toluene observed in this study.

In summary, our results provide, for the first time, evidence that exposure of *S. schenckii* to soil contaminant concentrations of toluene can induce fungal modifications that could contribute to virulence. New studies are being conducted to elucidate the biochemical and molecular changes at the cellular level in response to exposure to toluene and other related xenobiotics, as well as=other changes induced in the immune system. These results support the hypothesis that the environment has a direct influence on the virulence of the fungus and constitutes an attractive way to explain, at least in part, some cases of sporotrichosis outbreaks in highly polluted regions.

## Acknowledgments

This study was financed in part by the Coordenação de Aperfeiçoamento de Pessoal de Nível Superior - Brasil (CAPES) - Finance Code 001

## References

Agency for Toxic Substances and Disease Registry. ATSDR’s Substance Priority List. (2017).

Alegranci, P., de Abreu Ribeiro LC, Ferreira LS, Negrini Tde C, Maia DC, Tansini A, Gonçalves AC, Placeres MC, Carlos IZ. (2013). The predominance of alternatively activated macrophages following challenge with cell wall peptide-polysaccharide after prior infection with Sporothrix schenckii. Mycopathologia.;176(1-2):57-65. doi:10.1007/s11046-013-9663-y.

Almeida-Paes, R., Figueiredo-Carvalho, M. H. G., Brito-Santos, F., Almeida-Silva, F., Oliveira, M. M. E., & Zancopé-Oliveira, R. M. (2016). Melanins protect Sporothrix brasiliensis and Sporothrix schenckii from the antifungal effects of terbinafine. PLoS ONE;11(3):e0152796.. https://doi.org/10.1371/journal.pone.0152796

Alviano, C.S., Farbiarz, S.R., De Souza, W., Angluster, J., Travassos, L.R. (1991) Characterization of Fonsecaea pedrosoi melanin. J Gen Microbiol.;137(4):837–44. https://doi.org/10.1099/00221287-137-4-837

Angelova, M. B., Pashova, S. B., Spasova, B. K., Vassilev, S. V., & Slokoska, L. S. (2005). Oxidative stress response of filamentous fungi induced by hydrogen peroxide and paraquat. Mycological Research. 109(Pt 2):150–8. https://doi.org/10.1017/S0953756204001352

Badali, H., Gueidan, C., Najafzadeh, M. J., Bonifaz, A., Gerrits van den Ende, A. H. G., & de Hoog, G. S. (2008). Biodiversity of the genus Cladophialophora. Studies in Mycology. 61:175–91. https://doi.org/10.3114/sim.2008.61.18

Barros, M. B. d L., Schubach, A. O., Schubach, T. M. P., Wanke, B., & Lambert-Passos, S. R. (2008). An epidemic of sporotrichosis in Rio de Janeiro, Brazil: Epidemiological aspects of a series of cases. Epidemiology and Infection. 136(9):1192–6. https://doi.org/10.1017/S0950268807009727

Barros, M. B. de L., de Almeida Paes, R., & Schubach, A. O. (2011). Sporothrix schenckii and Sporotrichosis. Clinical Microbiology Reviews. 24(4):633–54. https://doi.org/10.1128/CMR.00007-11

Batista-Duharte A., Martínez D.T., da Graça Sgarbi D.B., Carlos I.Z. (2015) Environmental Conditions and Fungal Pathogenicity. In: Zeppone Carlos I. (eds) Sporotrichosis. Springer, Cham. https://doi.org/10.1007/978-3-319-11912-0_4.

Batista-Duharte, A., Téllez-Martínez, D., Jellmayer, JA, Portuondo Fuentes DL, Campos Polesi M, Baviera A M, Zeppone Carlos I. (2018) Repeated Exposition to Mercury (II) Chloride Enhances Susceptibility to S. schenckii sensu stricto Infection in Mice. J Fungi (Basel).;4(2). pii: E64. doi:10.3390/jof40200.

Batista-Duharte, A., Téllez-Martínez D, Roberto de Andrade C, Portuondo DL, Jellmayer JA, Polesi MC, Carlos IZ. (2018) Sporothrix brasiliensis induces a more severe disease associated with sustained Th17 and regulatory T cells responses than Sporothrix schenckii sensu stricto in mice. Fungal Biol. Dec;122(12):1163-1170. doi:10.1016/j.funbio.2018.08.004

Blasi, B., Poyntner, C., Rudavsky, T., Prenafeta-Boldú, F. X., Hoog, S. De, Tafer, H., & Sterflinger, K. (2016). Pathogenic yet environmentally friendly? black fungal candidates for bioremediation of pollutants. Geomicrobiology Journal. 33(3-4):308–317. https://doi.org/10.1080/01490451.2015.1052118

Blasi, B., Tafer, H., Kustor, C., Poyntner, C., Lopandic, K., & Sterflinger, K. (2017). Genomic and transcriptomic analysis of the toluene degrading black yeast Cladophialophora immunda. Scientific Reports. 7(1):11436. https://doi.org/10.1038/s41598-017-11807-8

Borba, C. D. M., Silva, A. M. M., & Oliveira, P. C. De. (1992). Long-time survival and morphological stability of preserved *Sporothrix schenckii* strains. Mycoses. 35(7-8):185–8.

Carlos, I., & Batista-Duharte, A. (2015). *Sporotrichosis: An emergent disease*. Sporotrichosis: New Developments and Future Prospects. In: Zeppone Carlos I. (eds) Sporotrichosis. Springer, Cham. https://doi.org/10.1007/978-3-319-11912-0_1

Casadevall, A., & Pirofski, L. A. (2007). Accidental virulence, cryptic pathogenesis, martians, lost hosts, and the pathogenicity of environmental microbes. Eukaryotic Cell. 6(12):2169–74. https://doi.org/10.1128/EC.00308-07

Casadevall, A., Steenbergen, J. N., & Nosanchuk, J. D. (2003). “Ready made” virulence and “dual use” virulence factors in pathogenic environmental fungi - The Cryptococcus neoformans paradigm. Current Opinion in Microbiology. 6(4):332–7. https://doi.org/10.1016/S1369-5274(03)00082-1

Chakrabarti, A., Bonifaz, A., Gutierrez-Galhardo, M. C., Mochizuki, T., & Li, S. (2014). Global epidemiology of sporotrichosis. Medical Mycology. 53(1):3–14. https://doi.org/10.1093/mmy/myu062

de Capriles, C. H., Essayag, S. M., Lander, A., & Camacho, R. (1993). Experimental pathogenicity of Sporothrix schenckii preserved in water (Castellani). Mycopathologia. 122(3):129–33. https://doi.org/10.1007/BF01103472

de Souza Lacerda, C. M. Martins, E. M. do N., de Resende, M. A., & de Andrade, A. S. R. (2011). Gamma Radiation Effects on Sporothrix schenckii Yeast Cells. Mycopathologia, 171(6), 395–401. https://doi.org/10.1007/s11046-011-9395-9

Dixon, D. M., Salkin, I. F., Duncan, R. A., Hurd, N. J., Haines, J. H., Kemna, M. E., & Coles, F. B. (1991). Isolation and characterization of Sporothrix schenckii from clinical and environmental sources associated with the largest U.S. epidemic of sporotrichosis. Journal of Clinical Microbiology. 29(6):1106–13.

Eisenman, H.C., Casadevall, A. (2012) Synthesis and assembly of fungal melanin. Appl Microbiol Biotechnol.;93(3):931-40. doi:10.1007/s00253-011-3777-2.

Ferreira LS, Gonçalves AC, Portuondo DL, Maia DC, Placeres MC, Batista-Duharte A, et al. (2015). Optimal clearance of Sporothrix schenckii requires an intact Th17 response in a mouse model of systemic infection. Immunobiology.;220(8):985–92.

Fernandes KS, Neto EH, Brito MM, Silva JS, Cunha FQ, Barja-Fidalgo C. Detrimental role of endogenous nitric oxide in host defence against Sporothrix schenckii. Immunology. 2008;123(4):469–79.

Franzen, A.J., Cunha MM, Miranda K, Hentschel J, Plattner H, da Silva MB, Salgado CG, de Souza W, Rozental S. (2008). Ultrastructural characterization of melanosomes of the human pathogenic fungus Fonsecaea pedrosoi. J Struct Biol.; 162:75–84.

Franzen, A.J., de Souza W, Farina M, Alviano CS, Rozental S. (1999). Morphometric and densitometric study of the biogenesis of electron-dense granules in Fonsecaea pedrosoi. FEMS Microbiol Lett.; 173:395–402.

Garrison, R.G., Boyd KS, Kier AB, Wagner JE. (1979) Spontaneous feline sporotrichosis: a fine structural study. Mycopathologia.;69(1-2):57–62.

Garrison, R.G., Mariat F, Boyd KS, Fromentin H. (1977). Ultrastructural observations of an unusual osmiophilic body in the hyphae of Sporothrix schenckii and Ceratocystis stenoceras. Ann Microbiol (Paris).;128(3):319–37.

Gessler, N. N., Aver’yanov, A. A., & Belozerskaya, T. A. (2007). Reactive oxygen species in regulation of fungal development. Biochemistry (Moscow). 72(10):1091–109. https://doi.org/10.1134/S0006297907100070

Giraud, T., Gladieux, P., & Gavrilets, S. (2010). Linking the emergence of fungal plant diseases with ecological speciation. Trends in Ecology and Evolution. 25(7):387–95. https://doi.org/10.1016/j.tree.2010.03.006

Gonçalves, A.C., Maia DC, Ferreira LS, Monnazzi LG, Alegranci P, Placeres MC, Batista-Duharte A, Carlos IZ. (2015) Involvement of major components from Sporothrix schenckii cell wall in the caspase-1 activation, nitric oxide and cytokines production during experimental sporotrichosis. Mycopathologia.;179(1-2):21-30. doi:10.1007/s11046-014-9810-0

Gonçalves, A.C., Ferreira LS, Manente FA, de Faria CMQG, Polesi MC, de Andrade CR, Zamboni DS, Carlos IZ. (2017). The NLRP3 inflammasome contributes to host protection during Sporothrix schenckii infection. Immunology.;151(2):154-166. doi:10.1111/imm.12719.

Gremião, I. D. F., Miranda, L. H. M., Reis, E. G., Rodrigues, A. M., & Pereira, S. A. (2017). Zoonotic Epidemic of Sporotrichosis: Cat to Human Transmission. PLoS Pathogens, 13(1), 2–8. https://doi.org/10.1371/journal.ppat.1006077

Kuo, W.-Y., Huang, C.-H., Shih, C., & Jinn, T.-L. (2013). Cellular Extract Preparation for Superoxide Dismutase (SOD) Activity Assay. BIO-PROTOCOL. https://doi.org/10.21769/BioProtoc.811

Kusuhara, M. (2009). Sporotrichosis and dematiaceous fungal skin infections. Japanese Journal of Medical Mycology. 50(4):213–7. https://doi.org/10.3314/jjmm.50.213

Lopes-Bezerra, L. M., Walker, L. A., Niño-Vega, G., Mora-Montes, H. M., Neves, G. W. P., Villalobos-Duno, H., Barreto L, Garcia K, Franco B, Martínez-Álvarez J.A, Munro C.A., Gow, N. A. R. (2018). Cell walls of the dimorphic fungal pathogens Sporothrix schenckii and Sporothrix brasiliensis exhibit bilaminate structures and sloughing of extensive and intact layers. PLoS Neglected Tropical Diseases, 12(3), 1–25. https://doi.org/10.1371/journal.pntd.0006169

Madrid, I. M., Mattei, A. S., Soares, M. P., Nobre, M. de O., & Meireles, M. C. A. (2011). Ultrastructural study of the mycelial phase of clinical isolates of Sporothrix schenckii obtained from feline, canine and human cases of sporotrichosis. Brazilian Journal of Microbiology. 42(3):1147–50. https://doi.org/10.1590/S1517-83822011000300037

Maia, D.C., Sassa, M.F., Placeres, M.C., Carlos, I.Z., (2006). Influence of Th1/Th2 cytokines and nitric oxide in murine systemic infection induced by Sporothrix schenckii. Mycopathologia 161, 11–19.

Maia, D.C., Gonçalves, A.C., Ferreira, L.S., Manente, F.A., Portuondo, D.L., Vellosa, J.C., Polesi, M.C., Batista-Duharte, A., Carlos, I.Z. (2016). Response of Cytokines and Hydrogen Peroxide to Sporothrix schenckii Exoantigen in Systemic Experimental Infection. Mycopathologia.;181(3-4):207-15. doi:10.1007/s11046-015-9966-2.

Marimon, R., Cano J, Gené J, Sutton DA, Kawasaki M, Guarro J. (2007) Sporothrix brasiliensis, S. globosa, and S. mexicana, three new Sporothrix species of clinical interest. J Clin Microbiol.;45(10):3198-206. doi:10.1128/JCM.00808-07.

Mendoza, M., Alvarado, P., Díaz de Torres, E., Lucena, L., & de Albornoz, M. C. (2005). [Physiological comportment and in vivo sensitivity of Sporothrix schenckii isolates maintained for 18 years by two preservation methods]. Rev Iberoam Micol. 22(3):151–6.

Morris-Jones, R., Youngchim, S., Gomez, B. L., Aisen, P., Hay, R. J., Nosanchuk, J. D., … Hamilton, A. J. (2003). Synthesis of melanin-like pigments by Sporothrix schenckii in vitro and during mammalian infection. Infection and Immunity. 71(7):4026–33. https://doi.org/10.1128/IAI.71.7.4026-4033.2003

Niyomploy, P., Chantragan, S., Daranee, Ch., Nawaporn, V., Aphichart, K., Polkit, S. (2014). Superoxide dismutase isozyme detection using two-dimensional gel electrophoresis zymograms. Journal of Pharmaceutical and Biomedical Analysis.;(90)72–77.

Nosanchuk, J. D., Stark, R. E., & Casadevall, A. (2015). Fungal melanin: What do we know about structure? Frontiers in Microbiology. 6:1463. https://doi.org/10.3389/fmicb.2015.01463

Pasarell, L., & McGinnis, M. R. (1992). Viability of fungal cultures maintained at −70°C. Journal of Clinical Microbiology, 30(4), 1000–1004.

Pečiulytė, D. D.-V. V. (2010). Effect of long-term industrial pollution on microorganisms in soil of deciduous forests situated along a pollution gradient next to a fertilizer factory 3. Species diversity and community structure of soil fungi. Ekologija, 56, 132–143.

Pendreño Y, González-Párraga P, Conesa S, Martínez-Esparza M, Aguinaga A, Hernández JA, Argüelles JC. (2006). The cellular resistance against oxidative stress (H2O2) is independent of neutral trehalase (Ntc1p) activity in Candida albicans. FEMS Yeast Res.;6(1):57–62.

Prenafeta-Boldú, F. X., Summerbell, R., & Sybren De Hoog, G. (2006). Fungi growing on aromatic hydrocarbons: Biotechnology’s unexpected encounter with biohazard? FEMS Microbiology Reviews. 30(1):109–30. https://doi.org/10.1111/j.1574-6976.2005.00007.x

Ramírez-Quijas, M. D., Zazueta-Sandoval, R., Obregón-Herrera, A., López-Romero, E., & Cuéllar-Cruz, M. (2015). Effect of oxidative stress on cell wall morphology in four pathogenic Candida species. Mycological Progress. 14:8. https://doi.org/10.1007/s11557-015-1028-0

Ramírez-Soto, M. C., Aguilar-Ancori, E. G., Tirado-Sánchez, A., & Bonifaz, A. (2018). Ecological Determinants of Sporotrichosis Etiological Agents, 4(3). pii: E95. https://doi.org/10.3390/jof4030095

Renzoni, A., Andrey, D.O., Jousselin, A., Barras, C., Monod, A., Vaudaux P, Lew D, Kelley WL. (2011) Whole Genome Sequecing and Complete Genetic Analysis Reveals Novel Pathways to Glycopeptide Resistence in Staphylococcus aureus. PLoS ONE;6(6):e21577.

Rodrigues AM, de Hoog G, Zhang Y, de Camargo ZP. (2014). Emerging sporotrichosis is driven by clonal and recombinant Sporothrix species. Emerg Microbes Infect.;3(5):e32. https://doi.org/10.1038/emi.2014.33.

San-Blas G, Guanipa O, Moreno B, Pekerar S, San-Blas F. (1996). Cladosporium carrionii and Hormoconis resinae (C. resinae): cell wall and melanin studies. Curr Microbiol.; 32:11–16.

Shao, J., Shi, G. X., Wang, T. M., Wu, D. Q., & Wang, C. Z. (2016). Antiproliferation of berberine in combination with fluconazole from the perspectives of reactive oxygen species, ergosterol and drug efflux in a fluconazole-resistant Candida tropicalis isolate. Frontiers in Microbiology. 7:1516. https://doi.org/10.3389/fmicb.2016.01516.

Silva, M.B, Costa MM, Torres CC, Galhardo MC, Valle AC, Magalhães Mde A, Sabroza PC, Oliveira RM. Urban sporotrichosis: a neglected epidemic in Rio de Janeiro, Brazil. Cad Saude Publica. 2012;28(10):1867–80.

Téllez, M. D., Batista-Duharte, A., Portuondo, D., Quinello, C., Bonne-Hernández, R., & Carlos, I. Z. (2014). Sporothrix schenckii complex biology: Environment and fungal pathogenicity. Microbiology 160(Pt 11):2352–65. https://doi.org/10.1099/mic.0.081794-0

Torres-Guerrero, H., & Arenas-López, G. (1998). UV irradiation induced high frequency of colonial variants with altered morphology in Sporothrix schenckii. Medical Mycology. 36(2):81–7. https://doi.org/10.1046/j.1365-280X.1998.00125.x

Ulfig, K. (1994). The occurrence of keratinolytic fungi in the polluted environment of the Labedy District in Gliwice. Roczniki Panstwowego Zakladu Higieny. 45(4):337–46.

Ulfig, K., Terakowski, M., & Lukasik, W. (1996). A preliminary study on the occurrence of keratinolytic fungi in the street sweepings from Chorzów. Rocz Panstw Zakl Hig. 47(2):143–9.

Upadhyay, S., Xu, X., Lowry, D., Jackson, J. C., Roberson, R. W., & Lin, X. (2016). Subcellular Compartmentalization and Trafficking of the Biosynthetic Machinery for Fungal Melanin. Cell Reports. 14(11):2511–8. https://doi.org/10.1016/j.celrep.2016.02.059

Walker, C.A., Gomez BL, Mora-Montes HM, Mackenzie KS, Munro CA, Brown AJ, Gow NA, Kibbler CC, Odds FC. (2010). Melanin externalization in Candida albicans depends on cell wall chitin structures. Eukaryot Cell. 2010; 9:1329–1342.

Weydert, C. J., & Cullen, J. J. (2010). Measurement of superoxide dismutase, catalase and glutathione peroxidase in cultured cells and tissue. Nature Protocols. 5(1):51–66. https://doi.org/10.1038/nprot.2009.197

Wilson, M. E. (1995). Infectious diseases: An ecological perspective. BMJ. 311:1681–1684.https://doi.org/10.1136/bmj.311.7021.

